# Cytosolic sequestration of spatacsin by Protein Kinase A and 14-3-3 proteins

**DOI:** 10.1101/2020.09.09.289009

**Authors:** Susanna Cogo, James E. Tomkins, Nikoleta Vavouraki, Veronica Giusti, Federica Forcellato, Cinzia Franchin, Isabella Tessari, Giorgio Arrigoni, Laura Cendron, Claudia Manzoni, Laura Civiero, Patrick A. Lewis, Elisa Greggio

## Abstract

Mutations in *SPG11*, encoding spatacsin, constitute the major cause of autosomal recessive Hereditary Spastic Paraplegia (HSP) with thinning of the corpus callosum. Previous studies showed that spatacsin orchestrates cellular traffic events through the formation of a coat-like complex and its loss of function results in lysosomal and axonal transport impairments. However, the upstream mechanisms that regulate spatacsin trafficking are unknown. Here, using proteomics and CRISPR/Cas9-mediated tagging of endogenous spatacsin, we identified a subset of 14-3-3 proteins as physiological interactors of spatacsin. The interaction is modulated by Protein Kinase A (PKA)-dependent phosphorylation of spatacsin at Ser1955, which initiates spatacsin trafficking from the plasma membrane to the intracellular space. Our study provides novel insight in understanding spatacsin physio-pathological roles with mechanistic dissection of its associated pathways.

## Introduction

Loss of function mutations in *SPG11*, encoding spatacsin, constitute the major cause of autosomal recessive Hereditary Spastic Paraplegia with thinning of the corpus callosum (ARHSP-TCC). Although being largely heterogeneous, all HSPs are unified by the degeneration of distal axons in the corticospinal tracts, clinically resulting in slowly progressive spasticity primarily affecting the lower limbs (Blackstone, 2020; Stevanin et al., 2008). Genetically, >80 causative *loci* and over 60 genes have been identified and Mendelian modes of inheritance together with mitochondrial mutations have been described to date (Blackstone, 2018). Despite the large amount of genetic contributors, HSP genes have been associated with a relatively small number of cellular functions related to organelle biogenesis and membrane trafficking, lipid and mitochondria metabolism, development and myelination, as well as axonal transport and autophagy (Darios et al., 2020; Blackstone, 2018; Lo Giudice et al., 2014). Spatacsin is a 270kDa protein, highly expressed in the central nervous system, particularly in the motor cortex, spinal cord, hippocampus, cerebellum, dentate nucleus and pons (Murmu et al., 2011). Despite its association with multiple neurodegenerative disorders, including HSP, juvenile Parkinson’s disease, Charcot-Marie-Tooth disease, amyotrophic lateral sclerosis and Kjellin’s syndrome (Patto & O’kane, 2020), very little is yet known about the physiological role(s) of spatacsin and the consequences of its loss of function. Previous studies suggested an involvement of spatacsin in cellular traffic events, through the formation of a coat-like complex in association with two other HSP-linked proteins: spastizin (encoded by *SPG15*) and the Adaptor Protein 5 (AP5) complex subunit zeta-1 (*SPG48*) (Hirst et al., 2013). Accordingly, patient-derived fibroblasts and knockout (KO) animal models for *SPG11* and *SPG15* or their homologs display overlapping cellular phenotypes, with the deposition of large membrane-surrounded materials that are positive for LAMP1 and, in neurons, accumulate prior to neuronal loss (Khundadze et al., 2019, 2013; Renvoisé et al., 2014; Varga et al., 2015). Recently, Darios and collaborators generated a *Spg11*^*-/-*^ mouse model that fully recapitulates the symptoms of HSP. At the molecular level, the mouse shows lysosomal dysfunction due to an impairment in lipid, and specifically ganglioside, clearance (Boutry et al., 2018; Branchu et al., 2017). Of interest, defects in both spatacsin and spastizin have been associated with an impairment of the autophagic-lysosome reformation (ALR) process with the inhibition of tubule formation from late endosomes/lysosomes, a mechanism that is suggested to be AP5-independent (Boutry et al., 2019; Vantaggiato et al., 2019; Chang et al., 2014). Notably, the impairment in lysosome tubulation observed in absence of spatacsin has been associated with deficits in the clearance of lysosomal cholesterol and consequent alterations in its subcellular distribution (Branchu et al., 2017). In addition, the loss of spatacsin has been linked with axonal pathologies, altered vesicular transport and reduced neurite complexity (Pérez-Brangulí et al., 2014), possibly dependent on a deregulation of the GSK3β/β-catenin signalling pathway (Pozner et al., 2018). Thus, spatacsin likely plays additional roles in axonal maintenance and cargo trafficking. Nevertheless, currently available data on the subcellular localisation of spatacsin are conflicting, with initial studies reporting colocalisation with the microtubules, endoplasmic reticulum (ER) and vesicles involved in protein trafficking (Murmu et al., 2011), and more recent data suggesting spatacsin resides in the endolysosomal compartment, and in neurons is found in both axonal and dendritic processes (Hirst et al., 2013; Pérez-Brangulí et al., 2014). In addition, subcellular localisation data from The Human Protein ATLAS (https://www.proteinatlas.org/ENSG00000104133-SPG11/subcellular) suggest that the protein is distributed between the cytosol and the plasma membrane, in support of a highly dynamic sub-compartmentalisation. A more detailed understanding of spatacsin pathophysiology has so far been challenged by the lack of sensitive tools for spatacsin detection and the poorly characterised spatacsin interactome. The recent development of BioPlex 3.0, a proteome-scale, cell-line specific interaction network, has resulted in an expansion of the list of spatacsin candidate binding partners - boosting the opportunities to further explore spatacsin-related cellular pathways (Huttlin et al., 2020). Here, we adopted an analogous approach and performed a high-throughput affinity purification (AP) coupled to mass-spectrometry (MS) analysis of 3xFlag-tagged spatacsin immunopurified from HeLa cells and incubated with a murine brain lysate, to identify novel interactors. Next, we assessed these novel hits alongside literature-derived data obtained from the powerful Protein Interaction Network Online Tool (PINOT – (Tomkins et al., 2020)) and the BioPlex 3.0 dataset. After prioritisation of the candidate hits, we functionally validated the interaction between spatacsin and a subset of 14-3-3 chaperones *in vitro* and in cells. 14-3-3 proteins are molecular shuttles, which regulate localisation, activity and stability of their binding partners by interacting with specific phospho-motifs (Giusto et al., 2021; Sluchanko & Gusev, 2017). 14-3-3 proteins are highly expressed in the brain during development and have been linked to multiple neurodegenerative processes (Civiero et al., 2017; Cornell & Toyo-oka, 2017; Kaplan et al., 2017; McFerrin et al., 2017). We identified a 14-3-3 binding site in spatacsin, and demonstrated that the interaction is modulated through phosphorylation of spatacsin at Ser1955 by cyclic AMP (cAMP)-dependent Protein Kinase (PKA). Supporting the *in silico* prediction that spatacsin is potentially a transmembrane protein, we observed that PKA activation rapidly induces spatacsin translocation from the plasma membrane to the intracellular space. Our study provides novel insight into a developing understanding of spatacsin physio-pathological roles with mechanistic dissection of spatacsin associated pathways.

## Results

### AP-MS screening of the spatacsin interactome nominates 14-3-3 proteins as candidate binding partners

To expand the current knowledge on potential spatacsin related functional pathways, we interrogated its interactome, by performing an unbiased, high-throughput screen of candidate protein-protein interaction (PPI) partners *via* affinity purification (AP) coupled with tandem mass spectrometry (MS/MS). After immunopurifying 3xFlag-tagged spatacsin or GFP as a negative control, we performed a pull-down assay by incubating the bait proteins with murine brain lysate. Afterwards, we submitted the proteins bound to either spatacsin or GFP for MS/MS analysis. Eighty-four proteins were considered positive hits in terms of potential spatacsin binding partners (see Materials and Methods for processing of MS/MS data). These interactors were converted to their human orthologs using the DSRC Integrative Ortholog Prediction Tool (DIOPT – Hu et al., 2011), whereby direct and high confidence mouse-human orthologs were identified in 97.6% of cases; 2 orthologs were assigned with moderate confidence (Supplementary File 1). Next, these proteins were used to query the CRAPome repository, a dataset of common MS contaminants (Mellacheruvu et al., 2013), which resulted in discarding 9 proteins from further analyses due to their prevalence as known contaminants in MS experiments (Supplementary File 2). The retained interactors were visualised in a PPI network, together with the extent of curated PPI literature for human spatacsin, obtained from PINOT (7 interactors), and BioPlex 3.0 (a further 38 interactors) (Figure 1A). Data from PINOT and BioPlex 3.0 were referred to collectively as literature-derived PPI data. The novel MS-derived spatacsin interactors were further analysed in terms of subcellular localisation, based on Gene Ontology Cellular Component (GO CC) classification. GO CCs were grouped into semantic classes, based on semantic similarity, which were then grouped into location blocks and categories (Supplementary File 3). We further investigated the semantic composition of the GO CC terms associated with the spatacsin interactome by grouping terms related to neuronal structures or lysosomes into two additional location groups named “Neuron-related terms” and “Lysosomes”. This analysis showed that the localisation of spatacsin has been associated with membranes, cytosol, nucleus, other organelles, vesicles, and cellular projections, and that spatacsin had GO CC terms related to neuronal structures and lysosomes (Figure 1B). Of note, 34 out of 75 of spatacsin interactors were associated with neuron related GO CC terms (45.3%), 6 of which were 14-3-3 proteins, namely the isoforms beta, epsilon, gamma, eta, theta, and zeta (6/34, 17.6%).

**Figure 1.**
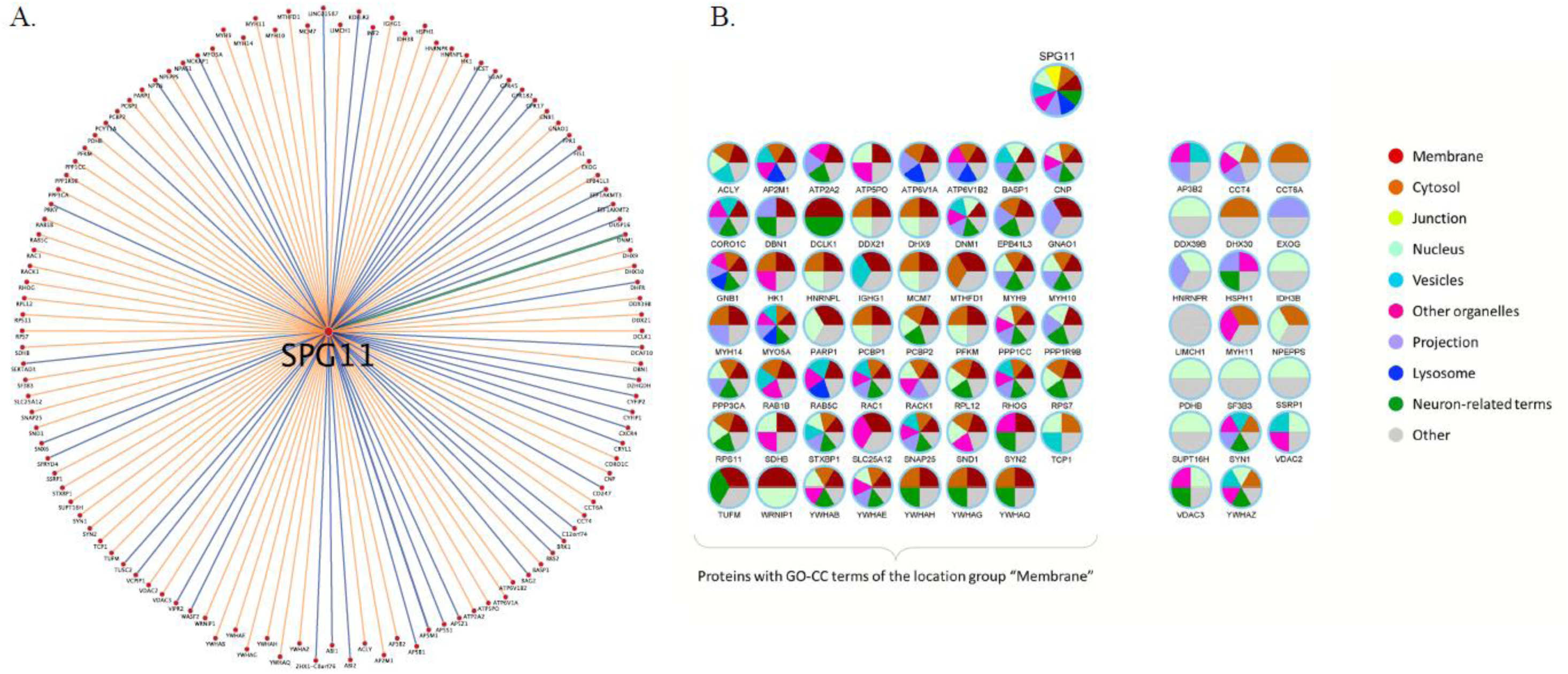
The spatacsin interactome. (A) A protein interaction network of spatacsin illustrating novel protein interactors identified by mass spectrometry in this study (orange edges), literature-derived protein interactors from PINOT and BioPlex 3.0 (blue edges), and an interactor common to both our novel dataset and literature-derived data (green edge). The edge thickness positively correlates with protein interaction confidence based on distinct method detection strategies used and the number of specific documented reports. (B) The subcellular localisation of the interactors resulted from our AP-MS analysis and of spatacsin is visualised, with each colour corresponding to a different location category. The collection of subcellular location data in the form of Gene Ontology cellular component terms was performed through AmiGO and the grouping of each term into location categories with an in-house grouping protocol.

### Spatacsin interacts with a subset of 14-3-3 proteins

Based on MS screening and functional prioritisation of spatacsin candidate binding partners, we decided to pursue validation of the interaction between spatacsin and the 14-3-3 protein family. 14-3-3s are a group of seven adaptors that participate in multiple cellular functions, assisting the localisation, activity and/or stability of their binding partners (Civiero et al., 2017; Sluchanko & Gusev, 2017). Relevant to spatacsin biology, 14-3-3s are highly expressed in the brain and are well-known to regulate the subcellular localisation of their client proteins through interaction with phosphorylated motifs (Sluchanko & Gusev, 2017; Foote & Zhou, 2012). Interestingly, six out of the seven 14-3-3 isoforms were pulled down during our AP-MS experiment (Figure 1A). We initially confirmed the interaction between 3xFlag-tagged spatacsin overexpressed in HEK293T cells and endogenous 14-3-3s, which were detected through a pan antibody designed against a common epitope shared by all the isoforms (Figure 2A). When assessing the subcellular localisation of 3xFlag-spatacsin in basal conditions, we found the protein to follow a punctate pattern with sporadic co-localisation with the ER (Figure 2B). Of note, a pool of spatacsin is present at the cell periphery in protrusion-like structures (Figure 2B), in agreement with the GO CC enrichment of spatacsin interactors at cellular projections (Figure 1B). Moreover, spatacsin staining partially colocalises with 14-3-3 proteins, supporting the notion that, under unstimulated conditions, a pool of 14-3-3s is compartmentalised with spatacsin (Figure 2C). At this point, in order to determine which 14-3-3 isoforms bind spatacsin with higher affinity, we purified *via* immobilised-metal affinity chromatography (IMAC) all seven 6xHis-tagged 14-3-3 family members and incubated them with affinity-purified 3xFlag-spatacsin. We observed that spatacsin can bind all 14-3-3 isoforms, with the only exception of σ-14-3-3, confirming the MS results, and preferentially binds isoforms η and γ, suggestive of a certain degree of specificity (Figure 2D).

**Figure 2.**
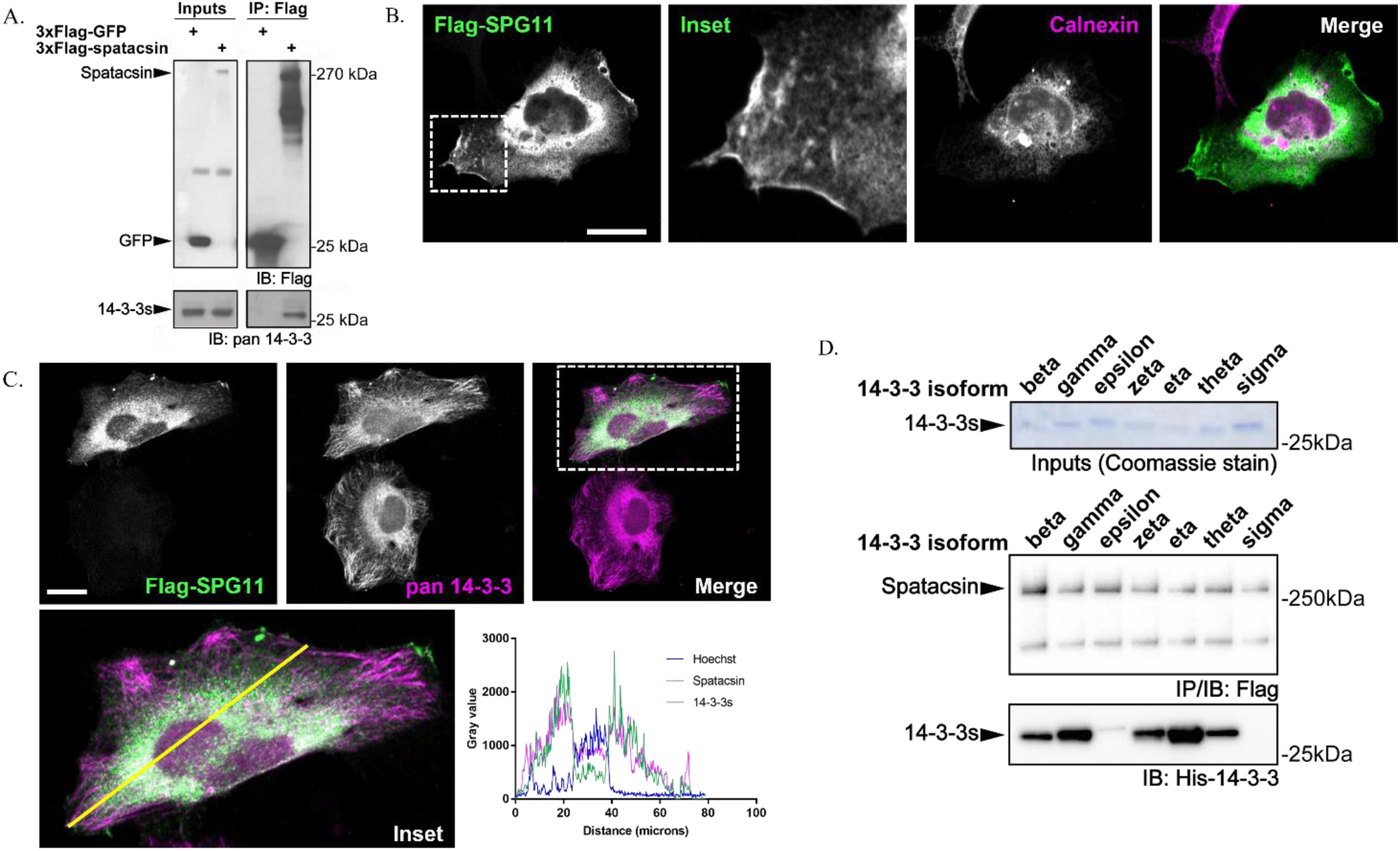
Spatacsin interacts with a subset of 14-3-3s in cells. (A) 3xFlag-spatacsin overexpressed in HEK293T cells pulls down endogenous 14-3-3 proteins. (B) Immunocytochemical evaluation of 3xFlag-spatacsin subcellular compartmentalisation in HeLa cells reveals predominant perinuclear localisation, partially overlapping with the ER marker calnexin. Of note, a pool of spatacsin is located at the cell periphery in protrusions-like structures. (C) Co-staining of 3xFlag-spatacsin and 14-3-3 proteins suggests that spatacsin only overlaps with a subset of 14-3-3s, as confirmed *via* pull-down assays between 3xFlag-spatacsin and recombinant 14-3-3 isoforms purified by IMAC (D). Scale bars=20µm.

### Phosphorylation at Ser1955 is key for spatacsin binding to 14-3-3 proteins

Considering that 14-3-3 proteins typically interact with their partners at specific phospho-sites, commonly phospho-serines, through recognition of conserved consensus sequences (Civiero et al., 2017; Sluchanko & Gusev, 2017), we interrogated the 14-3-3-Pred tool for the prediction of candidate 14-3-3 binding sites in spatacsin (http://www.compbio.dundee.ac.uk/1433pred – Madeira et al., 2015) and the PhosphoSitePlus® database (Hornbeck et al., 2015), which curates information on protein post-translational modifications, including phosphorylation. In parallel, we used phospho-peptide enrichment coupled with liquid-chromatography (LC)-MS/MS analysis on purified 3xFlag-spatacsin, to identify putative phosphorylated peptides. Figure 3A summarises the results obtained by the combination of *in silico* and experimental data. Among the seven candidate 14-3-3 binding sites predicted by 14-3-3-Pred, only Ser1955 (highlighted in orange) (i) was previously reported to be a phosphorylated residue in cells (119 references of high-throughput studies mapped in PhosphoSitePlus - https://www.phosphosite.org/homeAction) and (ii) was identified by our phospho-peptide enrichment. To validate Ser1955 as a binding site for 14-3-3 proteins, we generated the phospho-deficient S1955A (SA) mutant by site direct mutagenesis. We then performed pull-down assays between recombinant 3xFlag-spatacsin and 6xHis-14-3-3 isoforms to compare the binding affinity of wild-type (WT) versus mutant spatacsin for 14-3-3 proteins. Our results indicate that loss of phosphorylation on Ser1955 decreases 14-3-3s binding to spatacsin by ∼ 50% (Figure 3B).

**Figure 3.**
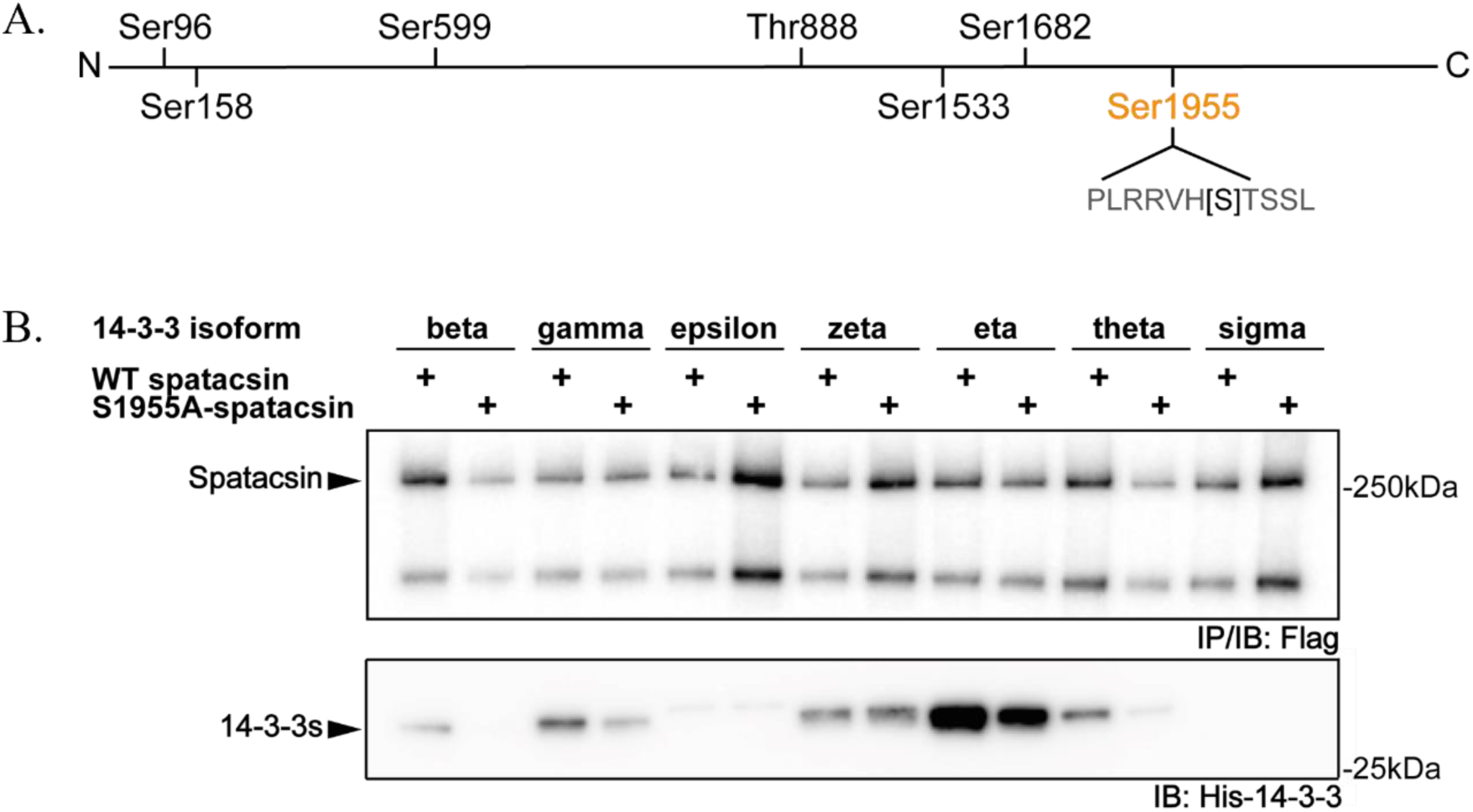
Phospho-Ser1955 is key for 14-3-3 binding to spatacsin. (A) Schematic of the putative 14-3-3 binding sites predicted by 14-3-3-Pred. Ser1955, highlighted in orange, was also reported to be a phosphorylated residue by PhosphoSitePlus®, and confirmed by phospho-peptide enrichment experiments performed in this work. (B) Binding affinity of 14-3-3s for phosphorylated spatacsin-Ser1955 was confirmed through generation of phospho-deficient S1955A-mutant by site-directed mutagenesis. 3x-Flag-spatacsin WT or S1955A were transfected in HeLa cells, immunoprecipitated with anti-Flag resin and incubated with equal amounts of recombinant 14-3-3s (1µM). The non-phosphorylatable mutant displays ∼ 50% reduction in binding as compared to the WT counterpart.

### PKA phosphorylates spatacsin at Ser1955

So far, the upstream mechanisms orchestrating spatacsin trafficking and compartmentalisation are poorly characterised. Based on the conserved mechanism through which 14-3-3s bind spatacsin *via* a phospho-motif, we employed the NetPhos 3.1 Server (Blom et al., 2004, 1999) to seek for the putative upstream kinase(s) responsible for phosphorylation of spatacsin at Ser1955. The results of the prediction, depicted in Figure 4A, strongly suggested PKA to be the upstream kinase phosphorylating spatacsin at this residue. At this point, we adopted a combination of cell biology and pharmacological approaches to validate the signalling cascade. Due to the lack of sensitive and selective antibodies against spatacsin, we generated monoclonal CRISPR/Cas9 genome-edited HeLa cell lines by inserting a 3xFlag-2xStrep tag at the N-terminus of the *SPG11* sequence. We selected HeLa cells as they represent the optimal compromise between ease of manipulation and suitability for the study of signalling and traffic pathways. To increase the likelihood of correct genome-editing, we designed two independent guide RNAs for the targeting of Cas9 in the proximity of the *SPG11* ATG start codon. A schematic of the experimental approach, together with the validation of CRISPR/Cas9-edited lines, are shown in Supplementary Figure 1A-C. As predicted by the design platform, sgRNA2 was more efficient, and clone 3 was selected for the following experiments (Supplementary Figure 1C). We subsequently immunopurified endogenous spatacsin from this cell line, taking advantage of the 3xFlag tag, in absence or presence of PKA pharmacological activation achieved with a combination of Forskolin and Isobutylmethylxanthine (IBMX), and monitored the binding of recombinant η-14-3-3 to spatacsin. Our data revealed that PKA activation significantly increases η-14-3-3 binding to spatacsin (Figure 4B), confirming the prediction that PKA phosphorylates spatacsin on Ser1955. In parallel, we took advantage of the CRISPR/Cas9-tagged spatacsin lines to evaluate the interaction between endogenous spatacsin and endogenous 14-3-3s in absence or presence of PKA activation by means of proximity ligation assay (PLA). As reported in Figure 4C, this confirmed the formation of the spatacsin/14-3-3 complex at the endogenous level, and indicates that heightened PKA activity increases the affinity of the spatacsin/14-3-3 complex.

**Figure 4.**
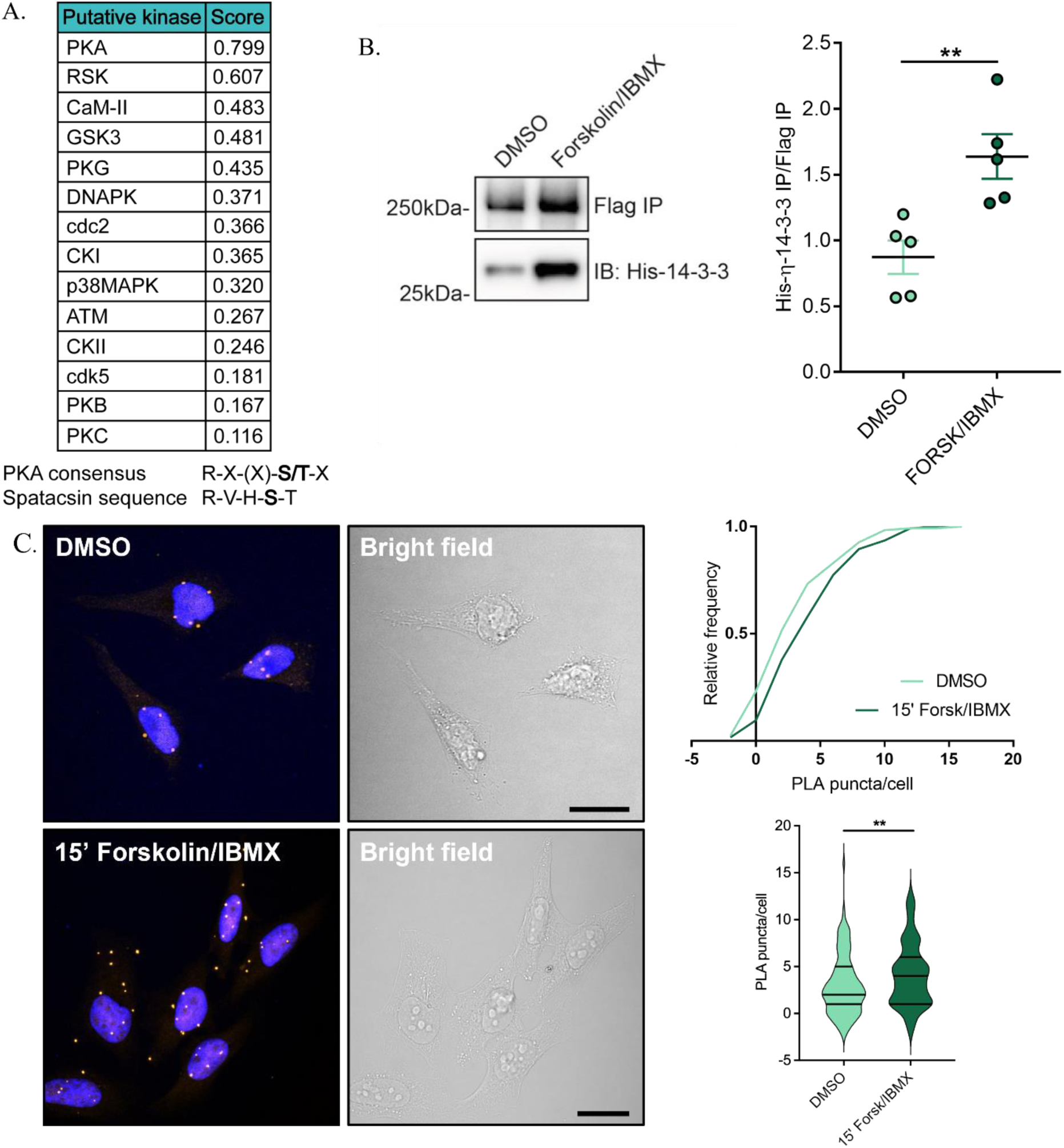
Ser1955 on spatacsin is phosphorylated by PKA. (A) Summary of the putative kinases phosphorylating spatacsin on Ser1955, with relative confidence score, as obtained from NetPhos 3.1. The PKA consensus sequence is also indicated and aligned against the spatacsin motif containing Ser1955. (B) Endogenous spatacsin was pulled down from genome-edited monoclonal lines taking advantage of the 3xFlag tag sequence inserted at the endogenous locus. Fifteen-minute treatment with Forskolin/IBMX increases binding affinity for η-14-3-3, suggesting that phosphorylation by PKA on Ser1955 is the signal regulating interaction (n=5, mean ± SEM, Unpaired t-test, ** p<0.01). (C) PLA experiments confirmed (i) interaction between spatacsin and 14-3-3s at the endogenous level and (ii) increased binding in the presence of PKA activation. PLA reactions were performed between rabbit α-Flag and mouse α-pan-14-3-3 antibodies. Average background signal detected in naïve cells, employed as a negative control for the reaction, was subtracted to all measurements; ∼125 cells counted/condition. Unpaired t-test, ^**^ p<0.01.

### PKA-mediated phosphorylation initiates spatacsin traffic from the plasma membrane

We next wanted to evaluate the functional consequence of PKA phosphorylation on spatacsin. PKA, the main cellular effector of cAMP, orchestrates several signalling pathways through phosphorylation of multiple substrates (Patra & Brady, 2018; Sassone-Corsi, 2012). In neurons, PKA produces a range of fast and dynamic responses at the plasma membrane, cytosol and the nucleus (within seconds at the membrane level and several minutes at the cytosol and nucleus levels), according to a tight spatiotemporal regulation of cAMP levels (Gervasi et al., 2007). In contrast, immortalised cell lines respond to PKA activation with slightly longer timescales (Hamaguchi et al., 2015; Namkoong et al., 2009). Previous evidence from bioinformatic studies suggests that spatacsin is a transmembrane protein (Stevanin et al., 2007). To further explore this, we utilised the structural prediction tools, PSIPRED and Phobius (http://bioinf.cs.ucl.ac.uk/psipred/ – Buchan & Jones, 2019; https://phobius.sbc.su.se/), which supported previous suggestions that spatacsin is a transmembrane protein with at least two transmembrane domains. Because a proportion of spatacsin appears to cluster in cellular protrusions consistent with a localisation at the plasma membrane (Figure 2B), we hypothesised that PKA-mediated phosphorylation on spatacsin-Ser1955 could change the proportion of spatacsin within these cellular pools. To test this, we employed a cell-surface biotinylation assay to measure (i) membrane versus intracellular spatacsin fractions and (ii) the response to PKA activation over time, *via* monitoring the internalisation of 3xFlag-2xStrep-tagged endogenous spatacsin after 5, 15 and 90 minutes of Forskolin/IBMX treatment. Figure 5A,B shows that (i) spatacsin is recovered in the membrane fraction and (ii) PKA activation reduces the plasma membrane fraction of spatacsin within 15 minutes, suggesting its internalisation. In parallel, we wanted to confirm (i) that this process is mediated by a 14-3-3-dependent mechanism and (ii) that it is potentially relevant in a pathophysiological context. Therefore, we imaged the localisation of spatacsin in murine primary striatal astrocytes transfected with spatacsin-WT or S1955A, treated over time with Forskolin/IBMX. In agreement with the results from the biotinylation assay, the amount of spatacsin located at the cell protrusions is significantly reduced after 15 minutes of PKA activation. Importantly, localisation of spatacsin-S1955A is unchanged upon Forskolin/IBMX treatment supporting a mechanism whereby PKA-mediated phosphorylation of Ser1955 is the signal that induces spatacsin trafficking from the plasma membrane to the intracellular compartments (Figure 5C,D; Supplementary Figure 2).

**Figure 5.**
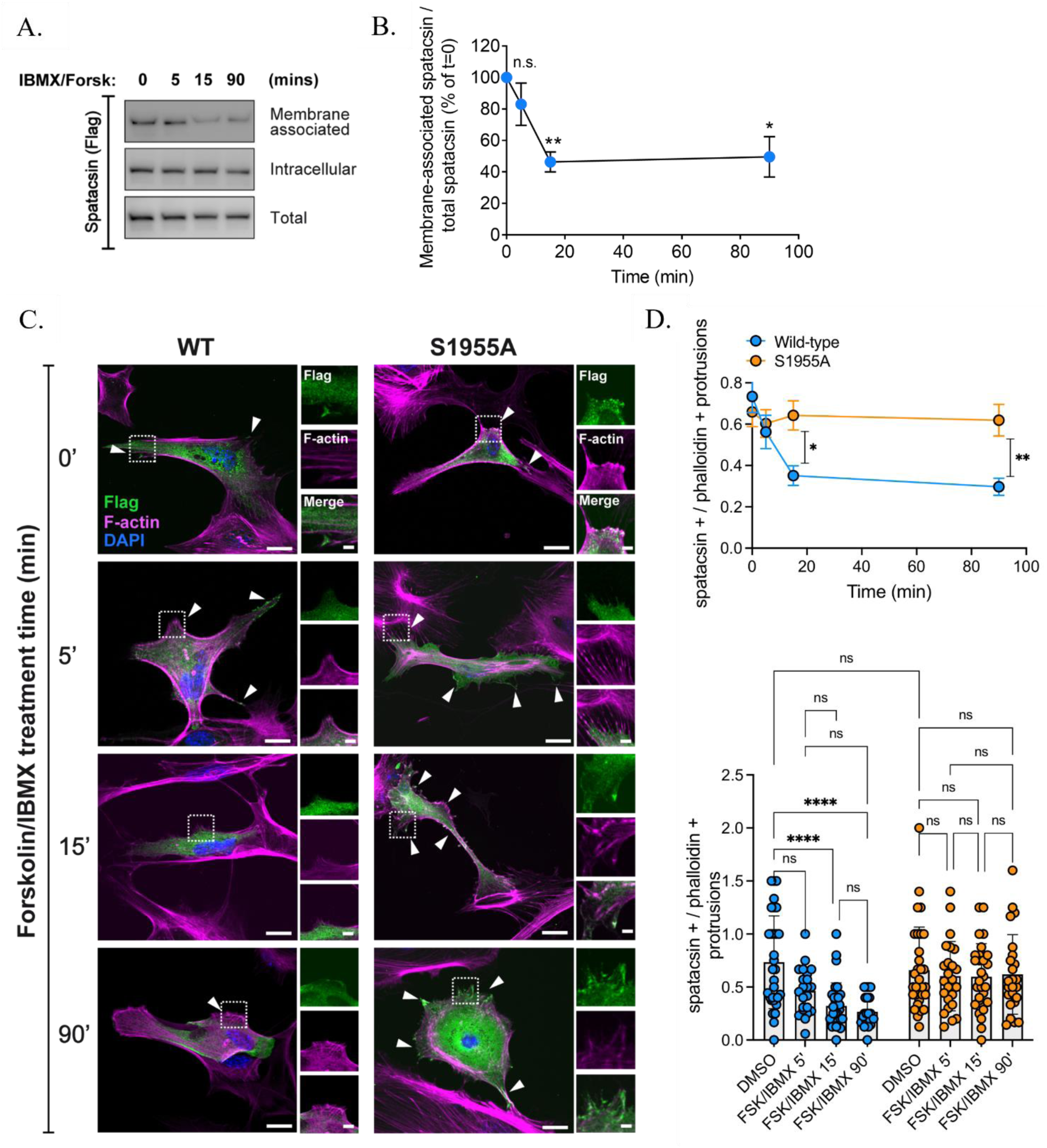
PKA-mediated phosphorylation initiates spatacsin traffic from the plasma membrane. (A) Cell surface biotinylation assay showing endogenous spatacsin internalisation upon 15 minutes treatment with Forskolin, suggesting that PKA activation is the stimulus to initiate spatacsin trafficking towards intracellular compartments. (B) Quantification of membrane-associated spatcsin over total protein (n=5, normalisation over time 0’, One-sample t-test against hyopetical value 100 (effect at t=0’ expressed as 100%); ^*^ p<0.05, ^**^ p<0.01. Normality tested with Shapiro-Wilk test). (C) Representative z-stack confocal images of primary striatal astrocytes trasfected with spatacsin-WT or S1955A under basal condition or treated with Forskolin/IBMX for 5, 15 and 90 minutes show that only the WT protein is able to respond to PKA activation. Insets show the colocalisation between spatacsin-WT or S1955A positive protrusions and phalloidin positive protrusions over time. Scale bar 20µm, insets 5 µm. (D) Quantification of data from (C), obtained from analysis of n=3 independent astrocyte cultures. Top: average values of spatacsin-positive protrusions normalised to phalloidin-positive protrusions over time. Bottom: distribution of values related to spatacsin-positive protrusions normalised to phalloidin-positive protrusions over time (two-way ANOVA, genotype ^****^ p<0.0001 F (3, 209) = 7.442, treatment ^***^ p<0.001, F (1, 209) = 14.91; multiple comparisons, ^*^ p<0.05, ^**^ p<0.01, ^***^ p<0.001, ^****^ p<0.0001). Note that in a small fraction of the spatacsin-positive protrusions no or poor co-localisation with phalloidin was detected.

## Discussion

Loss of function mutations in spatacsin represent the most common cause of ARHSP-TCC, and have been linked to multiple neurodegenerative disorders (Branchu et al., 2017). Previous studies demonstrated that lack of spatacsin leads to deficits in vesicle trafficking, axonal transport, ALR and lipid clearance (Boutry et al., 2019, 2018; Chang et al., 2014; Pérez-Brangulí et al., 2014; Hirst et al., 2013). However, detailed indications on spatacsin compartmentalisation and the mechanism of its regulation and recruitment are currently unavailable. We employed an unbiased, AP-MS approach and probed spatacsin as a bait against a whole brain sample, with the aim to identify binding partners of spatacsin in basal conditions. We then combined experimental data with cutting-edge bioinformatics approaches to guide the selection of candidate interactors for validation in a cellular system. We identified 75 putative novel interactors, which we prioritised according to their subcellular localisation. Intriguingly, we found the spatacsin interactome to be highly diverse in terms of subcellular compartmentalisation. Spatacsin itself was associated with multiple organelles, suggestive of a dynamic shuttling across compartments. The majority (n=54, 72%) of the hits identified from our MS analysis are associated with membranes, concordant with the idea of spatacsin acting as a coat-like protein able to mediate cellular traffic, possibly in connection with both the cytoskeleton and vesicles. Further corroborating this hypothesis, we identified specific sets of spatacsin interactors with established roles in regulating actin ruffling – Rho GTPases, Myosin isoforms, RAB proteins – and vesicle dynamics – e.g. members of the AP family – alongside members of multiple subcellular compartments (Figure 1B). Among all groups, we prioritised the hits that are most likely relevant in a neuronal context, based on the robust links between spatacsin and neurodegeneration, resulting in a list of 34 proteins. Strikingly, members of the 14-3-3 adaptor protein family represented the 17.6% of proteins with neuron-related GO CC terms. 14-3-3 proteins are abundantly expressed in the brain, with fundamental roles during development, and have been implicated in a variety of neurological disorders, based on genetic evidence, their relationship with relevant disease proteins and/or pathological findings (Foote & Zhou, 2012). Using phospho-peptide enrichment analysis and biochemical approaches, we identified and validated a phospho-residue at the C-terminus of spatacsin, namely Ser1955, as the main 14-3-3 binding site. Of note, depletion of Ser1955 phosphorylation, by means of a Ser-to-Ala phospho-deficient mutant, significantly reduced, although not completely abolished, this interaction. This can be explained by the fact that 14-3-3 proteins function predominantly as dimers, with each constituent monomer binding one phospho-peptide (Sluchanko & Gusev, 2012). Thus, additional 14-3-3 binding motifs are likely to be present within the spatacsin sequence, as also supported by *in silico* prediction from the 14-3-3-Pred tool (Figure 3A). Further supporting our results, an independent analysis performed with the Eukaryotic Linear Motifs (ELM – Kumar et al., 2020) software also suggests the presence of multiple 14-3-3 binding sites within spatacsin, including the RVHSTSSL peptide containing Ser1955. We additionally demonstrated Ser1955 to be phosphorylated by PKA. PKA activation is determined by an increase in cAMP levels, whereas specificity in the signalling is achieved through PKA-anchoring proteins (AKAPs), which position PKA in close proximity to effectors and substrates (Sassone-Corsi, 2012). Independent bioinformatic predictions performed by us and others suggest that spatacsin is a transmembrane protein, with multiple transmembrane domains (Figure 6). Based on the proposed roles of spatacsin in cellular trafficking, *via* AP5 and SPG15, on ALR, through binding with SPG15, and on our observation of a partial but consistent distribution of the protein at the level of the plasma membrane, we hypothesised that the interaction between spatacsin and 14-3-3s could play a pivotal role in the cellular localisation of spatacsin, with PKA-mediated phosphorylation acting as the triggering signal. We therefore demonstrated independently, through a cell-surface biotinylation assay on endogenously tagged HeLa cells and *via* immunofluorescence studies on HeLa cells and primary astrocytes transfected with WT and spatacsin-S1955A that PKA activation is capable of mediating spatacsin internalisation, and that this happens in a 14-3-3-dependent manner (Figure 5, Supplementary Figure 2). Future studies will be required to test the intriguing hypothesis that the actin cytoskeleton is assisting the internalisation, and to validate the molecular participants in the cascade. Therefore, given the interplay between *SPG11* and other spastic paraplegia genes, further mechanistic dissection of the PKA/spatacsin/14-3-3 cascade could shed light on the broader pathways that are impaired during disease. Gaining greater insight into the upstream signals that regulate this pathway, and whether these are AP5-dependent or independent events, will support the development of novel approaches to target their dysfunction in disease.

**Figure 6.**
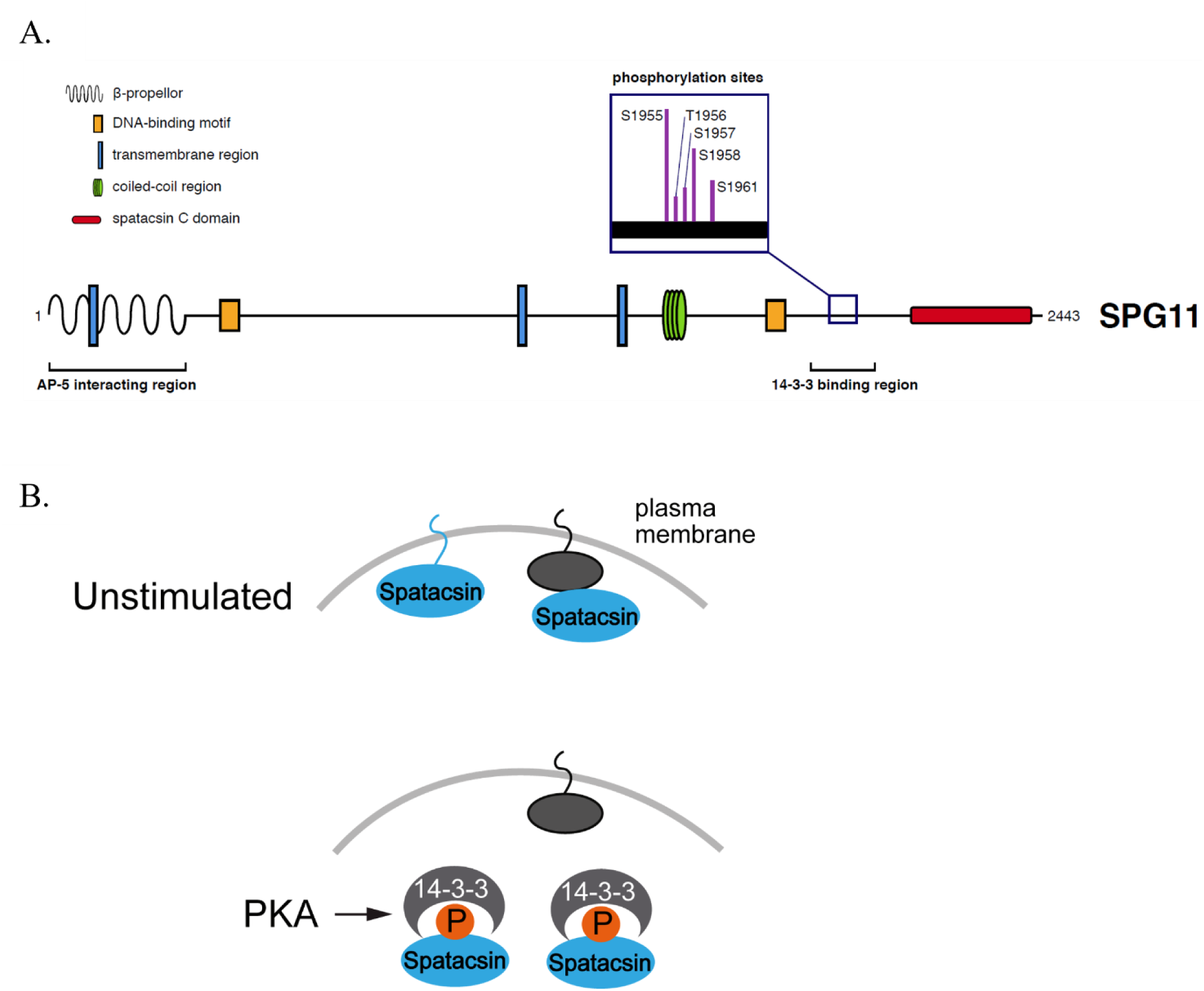
Cartoon representation of spatacsin domain organisation and current working model. (A) Canonical motifs for interactions with binding partners and transmembrane domains that have been predicted or identified by other groups (Hirst et al., 2013; Stevanin et al., 2007) and confirmed by *in silico* data from this study are summarised. The AP5-interacting region (Hirst et al., 2013) and the spatacsin C domain, which shares sequence and structure similarity with the Vps16 protein, identified in the study by Patto and O’Kane (Patto & O’kane, 2020) are also included, together with the phospho-peptide responsible for the binding of 14-3-3s identified in this study. (B) The findings from this study support previous data of spatacsin acting at the plasma membrane, although it remains unclear whether it is a transmembrane or a membrane-associated protein. Upon phosphorylation by PKA on spatacsin Ser1955, binding of 14-3-3 proteins to phospho-spatacsin induces its relocalisation to the intracellular space. Future studies will be required to obtain further mechanistic insight into the process.

## Experimental procedures

### Animals

Brain lysates were obtained from WT C57BL6J mice (The Jackson Laboratory). Housing and handling of mice were done in compliance with national guidelines. Animal procedures were approved by the Ethical Committee of the University of Padova and the Italian Ministry of Health (license 1041216-PR).

### Plasmids and reagents

The sequence of human *SPG11* was cloned into a pReceiver-M12 Expression Clone vector (GeneCopoeia). 3xFlag-GFP was used as a negative control in pull-down experiments. Generation of spatacsin-S1955A mutant was outsourced to VectorBuilder. Cloning of 6xHis-tagged 14-3-3 isoforms in a pET28a(+) vector for bacterial expression was performed as described previously (Civiero et al., 2017). For the generation of CRISPR/Cas9 edited lines, sgRNAs and the HDR template were synthesised by Sigma-Aldrich. sgRNAs were subsequently cloned into the pSpCas9(BB)-2A-Puro(PX459)V2.0 vector (Addgene). 2µg/ml puromycin (InvivoGen) was employed for the selection of positive clones. Treatment of cells with Forskolin and IBMX (both purchased from Sigma-Aldrich) was performed for the times indicated in figure captions at 30 and 100µM, respectively.

### Immortalised cell culture and transfection

HeLa and HEK293T cells purchased from ATCC were cultured in Dulbecco’s modified Eagle’s medium (DMEM, Life Technologies) supplemented with 10% foetal bovine serum (FBS, Life Technologies). Cell lines were maintained at 37°C in a 5% CO_2_ controlled atmosphere. 0.25% trypsin (Life Technologies), supplemented with 0.53mM EDTA, was employed to generate subcultures. Cells were transiently transfected with plasmid DNA using polyethylenimine (PEI, Polysciences), with a 1:3 DNA:PEI ratio. For pull-down assays, cells were plated onto 150mm dishes and transfected with 40μg of 3xFlag-spatacsin plasmid DNA. In the case of the 3xFlag-GFP control construct, which displays a higher efficiency in terms of both transfection and expression, only 10μg DNA were transfected, to obtain comparable purification yields. 48-72h after transfection, cells were harvested in RIPA buffer (Cell Signaling Technologies), supplemented with protease inhibitor cocktail (Sigma-Aldrich) and processed for the pull-down reactions.

### Primary astrocyte cell culture and transfection

Mouse primary striatal astrocytes were obtained from postnatal animals between day 1 and day 3 as previously described (Streubel-Gallasch et al., 2021). Briefly, brains were removed from the skull and striata were dissected under an optic microscope. After the dissection, Basal Medium Eagle (BME, Biowest), supplemented with 10% FBS (Life Technologies), 100 U/ml Penicillin + 100 µg/ml Streptomycin (Pen-Strep; Life Technologies), was added to the tissues. Striata were then sifted through a 70-μm cell strainer (Sarstedt) using a syringe plunger. The cell suspension was centrifuged (300 x g, 15 min) and the pellet was washed twice with 25 ml of supplemented medium. Cells were seeded at a density of 5×10^6^ cells/10 ml in cell culture T75 flasks. The culture medium was changed after 7 days and again after an additional 3-4 days. When cell confluency reached about 80%, microglia were detached by shaking the flask (800 rpm) for 2 h at room temperature (RT). After shaking, the medium containing microglia was replaced with fresh medium. Astrocytes were maintained in BME supplemented with 10% FBS and Pen-Strep at 37 °C in a controlled 5% CO_2_ atmosphere. After 14 days, astrocytes were used for the experiments. Transfection of primary striatal astrocytes was performed with 1µg 3xFlag-spatacsin WT or 3xFlag-spatacsin-S1955A plasmid using Lipofectamine 2000 (Thermo Scientific), following 1:3 plasmid to Lipofectamine ratio. After 48h, the experimental procedure was carried out.

### Generation of 3xFlag/2xStrep-spatacsin CRISPR/Cas9 edited monoclonal lines

Insertion of a 3xFlag/2xStrep tag at the N-terminus of endogenous spatacsin was performed using CRISPR/Cas9-mediated genome editing technology through homology-directed repair (HDR), following the protocol by Sharma et al., 2018. sgRNAs were designed taking advantage of the online platform Benchling (www.benchling.com) and scored based on previous publications (Doench et al., 2016; Hsu et al., 2013). Two sgRNAs were selected to increase the probability of success. As predicted by Benchling, sgRNA2 was found to be more specific and efficient in targeting *SPG11* (Supplementary Figure 1A). The 192bp tag sequence to be used as HDR template was obtained from Dalvai et al., 2015 and is flanked by two homology arms (HA), having a length of 600bp each and designed based on *SPG11* genomic sequence.

### Purification of recombinant 14-3-3 proteins

6xHis-14-3-3 isoforms were expressed in BL21(DE3) bacterial cells and purified in batch by IMAC with ProBond™ resin (Invitrogen). After elution with imidazole, buffer was exchanged to PBS with a PD10 desalting column (GE Healthcare). Proteins were quantified measuring UV absorption at 280nm and subsequently supplemented with 3mM dithiothreitol (DTT); proteins were quickly frozen and kept at − 80°C for long-term storage. 1:20 molar ratio between spatacsin and 14-3-3s was employed for pull-down experiments.

### Cell lysis, protein purification and pull-down assays

For the purification protocol, cell lysates were incubated on ice for 30 minutes and subsequently cleared by centrifugation at 20000g for 30 minutes at 4°C, after which the supernatants were incubated for 2 hours with 40μl of anti-Flag® M2 Affinity Gel (Sigma-Aldrich) at 4°C. After that, immunocomplexes were washed ten times in buffers with decreasing ionic strength and incubated with mouse brain lysates, or washed three times prior to proceeding to pull-downs with recombinant or endogenous 14-3-3s. The pull-down reactions were conducted overnight at 4°C in a binding buffer containing 20mM Tris-HCl, pH 7.5, 150mM NaCl, 1mM EDTA, 2.5mM sodium pyrophosphate, 1mM β-glycerophosphate, 1mM sodium orthovanadate and 1% NP-40. The following morning, resins were washed three times with the binding buffer and processed for Mass Spectrometry or Western Blot analysis.

### Cell-surface biotinylation assay

To evaluate the presence of spatacsin at the plasma membrane and its internalisation upon PKA activation, we performed the cell-surface biotinylation assay utilising the EZ-Link Sulfo-NHS-SS-Biotin (ThermoFisher Scientific) as per manufactures’ recommendations. Briefly, genome-edited cells were plated onto 6-well plates and subjected to treatment with PKA activators (Forskolin/IBMX) for 0, 5, 15 and 90 minutes. Cells were then transferred on ice and washed three times with ice-cold PBS pH 8.2, supplemented with 1mM MgCl_2_ and 0.1mM CaCl_2_ (PBS^2+^). Sulfo-Biotin was added for 30 minutes on ice at a concentration of 0.3mg/ml in PBS^2+^ in the dark. After three more washes in PBS^2+^, cells were harvested in RIPA buffer without EDTA, and biotinylated proteins were immunopurified through Streptavidin-Mag Sepharose™ (GE Healthcare). After washing of the beads, samples were processed for Western Blot. For each sample, input indicating the total amount of spatacsin and flow-through, representing the intracellular fraction of the protein, were also collected.

### Western Blot

15 to 30 µg of total protein samples were resolved on ExpressPlus™ PAGE 4–20% gels (GenScript), in MOPS running buffer, or 8% Tris-glycine polyacrylamide gels in SDS/Tris-glycine running buffer, according to the size-resolution required. The resolved proteins were transferred to polyvinylidenedifluoride (PVDF) membranes (Bio-Rad), through a Trans-Blot® Turbo™ Transfer System (Bio-Rad). PVDF membranes were subsequently blocked in Tris-buffered saline plus 0.1% Tween (TBS-T) and 5% non-fat milk for 1 hour at 4°C and then incubated over-night at 4°C with primary antibodies in TBS-T plus 5% non-fat milk. Membranes were then washed in TBS-T (3×10 minutes) at RT and subsequently incubated for 1 hour at RT with horseradish peroxidase (HRP)-conjugated α-mouse or α-rabbit IgG. Blots were then washed in TBS-T (4×10 min) at RT and rinsed in TBS-T, and immunoreactive proteins were visualised using Immobilon® Forte Western HRP Substrate (Merck Millipore). Densitometric analysis was carried out using the ImageJ 1.53c software. The antibodies used for Western Blot are as follows: mouse α-β-actin (A1978, Sigma-Aldrich, 1:20000), mouse α-pan 14-3-3 (sc-1657, Santa Cruz Biotechnology, 1:1000 to 1:10000), α-Flag® M2-HRP (A8592, Sigma-Aldrich, 1:20000), mouse α-His-HRP (A7058, Sigma-Aldrich, 1:10000).

### Protein digestion, phospho-peptide enrichment and LC-MS/MS analysis

Gel bands were subjected to in-gel digestion with sequencing-grade modified trypsin (Promega) as detailed in Belluzzi et al., 2016. A similar procedure was used for the digestion of phosphorylated proteins, but in this case LysC was used as protease. Briefly, gel bands were cut in small pieces, dehydrated with acetonitrile (ACN), and dried under vacuum. Reduction of cysteines was performed with freshly prepared 10mM DTT in 25mM Tris-HCl pH 8.5, at 56°C and for 1h. Alkylation was performed with 55mM iodoacetamide (Sigma Aldrich) in 25mM Tris-HCl pH 8.5 for 45mins at RT and in the dark. Gel pieces were washed several times with 25mM Tris-HCl pH 8.5 and ACN, dried under vacuum, and suspended LysC solution (Promega, 12.5ng/mL in 25mM Tris-HCl pH 8.5). Digestion was performed overnight at 37°C. Peptides were extracted with three changes of 50% ACN/0.1% formic acid (FA, Fluka). Samples were dried under vacuum and stored at − 20°C until the phosphopeptide enrichment procedure was performed. Phosphopeptide enrichment and LC-MS/MS analysis were performed as detailed in Belluzzi et al., 2016. MS analysis of the samples was performed with an LTQ-Orbitrap XL mass spectrometer (Thermo Fisher Scientific) coupled online with a nano-HPLC Ultimate 3000 (Dionex-Thermo Fisher Scientific). Raw data files were analysed with Proteome Discoverer software (version 1.4, Thermo Fisher Scientific) connected to a Mascot Server version 2.2.4 (Matrix Science, UK) against the Uniprot Mouse Database (version 2015.04.01). Trypsin or LysC were set as digesting enzymes with up to one missed-cleavage for protein identification and up to three missed-cleavages for phosphopeptides identification. Carbamidomethyl cysteine was set as fixed modification and methionine oxidation as variable modification in all cases. Phosphorylation of Ser/Thr/Tyr residues was set as variable modifications when the phosphopeptide analysis was performed. Peptide and fragment tolerance were 10 ppm and 0.6 Da, respectively. The algorithm Percolator was used to calculate False Discovery Rate (FDR) and PhosphoRS algorithm (Taus et al., 2011) was used to help in the assignment of the correct phosphorylation sites. Proteins were considered as correctly identified if at least 2 unique peptides per protein were sequenced with a q value <0.01. Quantification was achieved by integrating the area under the peaks for each identified peptide and then by averaging the values obtained for all peptides belonging to the same protein. A principle of maximum parsimony was applied to group proteins into protein families. The threshold for determining positive hits, in terms of spatacsin binding partners, was a minimum of 5 peptides matching the assigned protein and a minimum signal fold change above control of 5.

### Immunocytochemistry, PLA and confocal microscopy

For immunocytochemistry (ICC), cells were cultured onto 12mm glass coverslips in 12-well plates coated with poly-L-lysine (Sigma-Aldrich). Cells were fixed after 48h of transfection using 4% paraformaldehyde (PFA) and subsequently subjected to staining. The antibodies and dyes used for ICC are as follows: mouse α-pan 14-3-3 (sc-1657, Santa Cruz Biotechnology, 1:100), mouse α-Flag® M2 (F1804, Sigma-Aldrich, 1:100 to 1:400), polyHis-HRP (A7058, Sigma-Aldrich, 1:10000), rabbit α-DYKDDDDK tag – *alias* Flag – (14793S, CST, 1:100), Hoechst (33258, Invitrogen) and phalloidin-iFluor 647 (ab176759, abcam). For PLA, CRISPR/Cas9-edited lines were subjected to staining with the Duolink™ In Situ Detection Reagents Orange (DUO92007) and the combination of α-mouse and α-rabbit probes (DUO92002 & DUO92004) from Sigma-Aldrich, as per manufacturers’ recommendations. Proteins and complexes were visualised by confocal microscopy (Zeiss LSM700 and Leica SP5).

### Protein-protein interaction (PPI) networks and functional prioritisation of the data

Murine proteins identified by MS were converted to their respective human orthologs using the DSRC Integrative Ortholog Prediction Tool (DIOPT) version 8 (Hu et al., 2011); input species *Mus musculus*, output species *Homo sapiens*, with the option of all prediction tools selected to provide a broad meta-analysis. Ortholog assignments were determined based on the highest confidence predictions (Supplementary File 1). These orthologs were queried against the 716 datasets in the CRAPome repository, version 2.0 (Mellacheruvu et al., 2013) to identify likely contaminants in our MS analysis. Proteins present in >66.6% of the CRAPome datasets were discarded from further analyses (Supplementary File 2). Retained proteins were considered alongside literature-derived data, obtained from the Protein Interaction Network Online Tool version 1.1 (PINOT – Tomkins et al., 2020) and BioPlex 3.0 dataset (Huttlin et al., 2020), for network visualisation. PINOT was queried on 4th February 2022, with *Homo sapiens* and stringent filtering parameters selected. For the retained interactors of spatacsin from the AP-MS analysis, we collected the Gene Ontology Cellular Component (GO CC) terms through AmiGO v2 on 6^th^ August 2020, and for spatacsin on 21^st^ August 2020, using their HGNC approved name. The GO CC terms were grouped based on semantic similarity into semantic classes, which were further grouped into location blocks. Two additional location blocks were created, named “Neuron-related terms” and “Lysosome” in which we added any term that was related to neurons, or lysosomes, respectively. The amount of location blocks led to the need of further grouping into location categories to aid result interpretation. In addition, the categories that did not result from SPG11 were grouped in a new category “Other” for visualisation purposes. The existence of GO CC terms that belong in each category was then qualitatively visualised in Cytoscape using pie charts inside each node. Network and location categories visualisations were generated in Cytoscape v3.8.0.

### Bioinformatic predictions

Evaluation of putative transmembrane domains in spatacsin was performed using the PSIPRED predictor of secondary structure (Buchan & Jones, 2019) and Phobius, a combined transmembrane topology and signal peptide predictor developed by the Stockholm Bioinformatics Centre. Analysis of putative phosphorylation sites and the upstream kinases was conducted employing NetPhos 3.1 (ExPASy Bioinformatics Resource Portal). The Eukaryotic Linear Motif resource for Functional Sites in Proteins (ELM – Kumar et al., 2020; Puntervoll et al., 2003) and 14-3-3-Pred (http://www.compbio.dundee.ac.uk/1433pred – Madeira et al., 2015) were also interrogated for the prediction of 14-3-3 binding-motifs.

### Data analysis software

Measurement of relative band intensities in western blots, quantification of PLA positive signal and assessment of spatacsin internalisation were performed with ImageJ 1.53c software. For immunocytochemistry on primary astrocytes, images were acquired at 8-bit intensity resolution over 1024 × 1024 pixels, through Leica SP5 confocal microscope using a HC PL FLUOTAR x63/0.80 oil objective. Spatacsin-WT or 1955A and F-actin positive protrusions were manually counted using ImageJ 1.53c software. For each culture, eight independent fields per experiment were evaluated and reported. Imaging data were analysed blinded. Graphs and statistical analysis were realised using GraphPad PRISM 7.

