## Supplementary Figures for "Cytosolic sequestration of spatacsin by Protein Kinase A and 14-3-3 proteins"

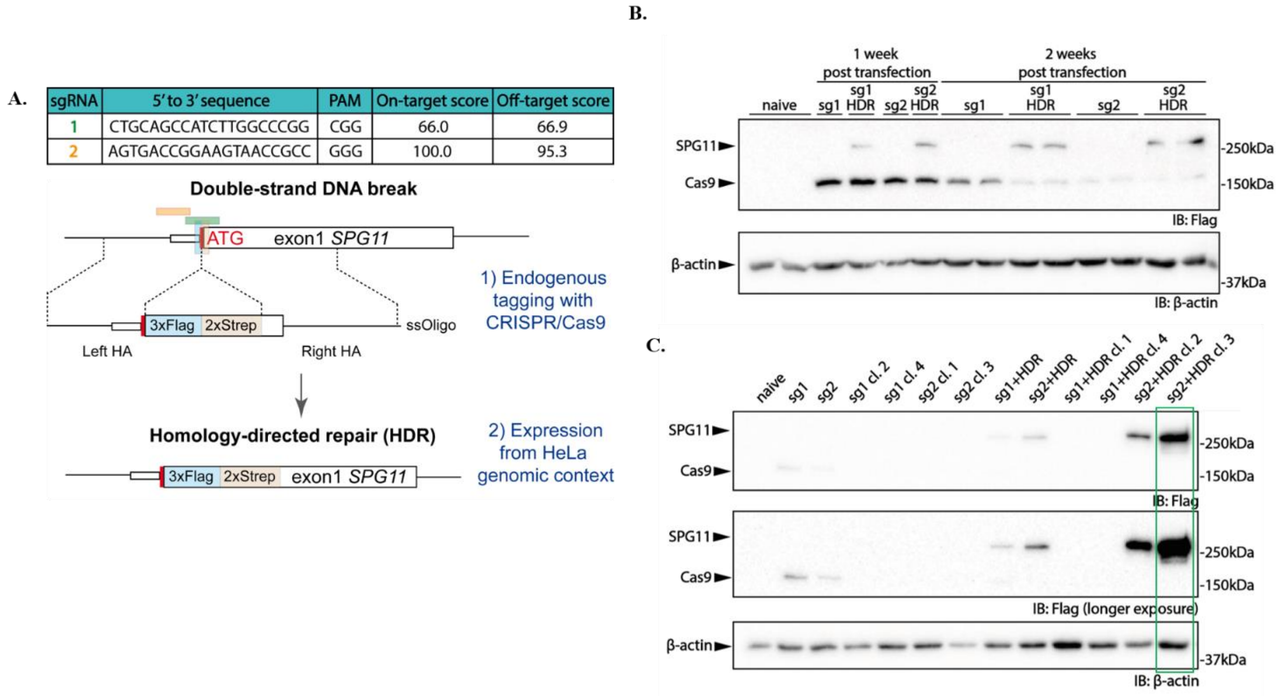


**Supplementary Figure 1 – Validation of the monoclonal cell lines generated through CRISPR/Cas9-mediated genome-editing.** (A) Schematic of the experimental setup, including sgRNA sequences and their scores, obtained through the Benchling online platform. Correct insertion of the tag was checked over time in the polyclonal (B) and monoclonal (C) cell lines through Western Blot against the Flag tag. Loss of the Cas9 plasmid was also monitored. sgRNA2 proved more efficient in correctly directing the HDR process. Clone 3 was selected for experimental procedures.

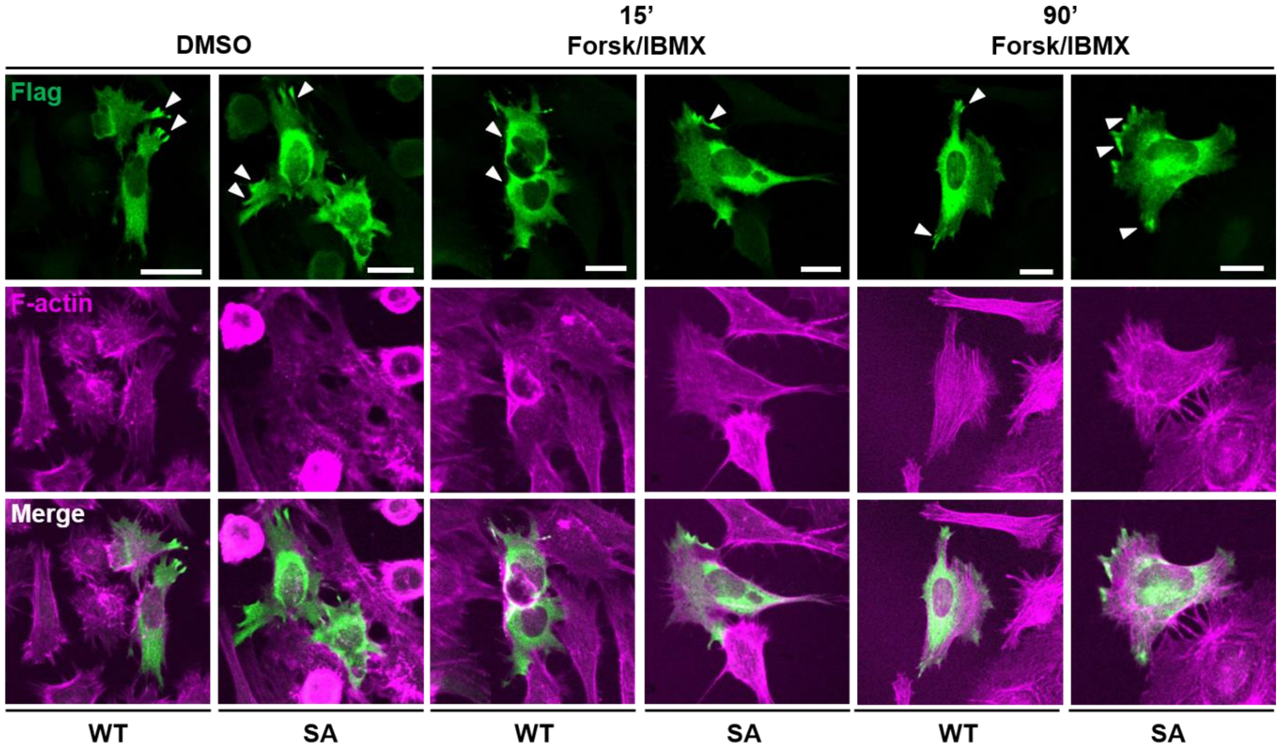


**Supplementary Figure 2 – PKA phosphorylates spatacsin on Ser1955 and induces its internalisation.** Representative images of HeLa cells transfected with spatacsin-WT or S1955A and treated with Forskolin/IBMX for 15 and 90 minutes show that only the WT protein is able to respond to PKA activation, suggesting that the mechanism is mediated by 14-3-3 binding. White arrowheads indicate spatacsin enrichment at protrusion-like structures, which is reduced after 15 minutes treatment with Forskolin/IBMX. Scale bar=20μm.
